# Engineering non-cytotoxic delivery of proteins by T cells via fusion to NPC2

**DOI:** 10.1101/2022.09.24.509028

**Authors:** Arash Saeedi, Irfan N. Bandey, Ankita Leekha, Mohsen Fathi, Ali Rezvan, Melisa J. Montalvo, Kwan-Ling Wu, Ritu Bohat, Weiyi Peng, Navin Varadarajan

## Abstract

Engineering cellular therapeutics by programming T cells has great potential in immunology. The primary mechanism employed by T cells for the specific transfer of proteins at the immunological synapse is via the lysosomal perforin pathway that facilitates the transfer of cytotoxic granzymes leading to apoptosis in target cells. Facilitating the delivery of non-cytotoxic proteins through perforin oligomers will dramatically expand the range of protein cargos that T cells can traffic to the target cells. Here, we have identified the intralysosomal protein, NPC2, as a chaperone that can facilitate the delivery of T-cell derived reporter proteins through perforin pores at the immunological synapse. Structural and biophysical considerations suggested that NPC2 could traverse through perforin pores and in vitro experiments confirmed the transport of purified NPC2 through perforin pores on cell membranes. To characterize the ability of NPC2 to facilitate the transfer of payloads in T cells, we constructed NPC2-mCherry fusion proteins in T cells. Using confocal microscopy and flow cytometry, we confirmed the colocalization of the NPC2 fused protein with lytic granules and the transfer of the fluorescent protein payload from T cells to target cells in co-culture experiments. The NPC2 fusion enabled the localization of mCherry to secretory lysosomes in mouse TCR CD8^+^ T cells and human CD4^+^ and CD8^+^ chimeric antigen receptor (CAR) T cells. These results illustrate that by using NPC2 as a molecular chaperone, the NPC2-perforin pathway can be exploited as a programmable molecular delivery system for cell-based therapies.

## Introduction

T cells are one of evolution’s cellular engineering masterpieces endowed with the ability to kill abnormal cells with high specificity. Recognition of mutated or foreign peptides restricted in the context of the appropriate major histocompatibility complex (MHC) molecule on the target cell results in the formation of the immunological synapse, a molecular conduit connecting the cells that can be used to deposit cytotoxic molecules leading to the apoptosis of the target cell. Activated CTLs are also capable of proliferation to make copies of themselves while also performing other functions like the secretion of messenger molecules like cytokines/chemokines. The ability of T cells to recognize external stimuli and respond by killing target cells and initiating self-proliferation has given birth to a new class of living drugs that is able to dynamically expand and contract in number matched to the disease burden. The engineering of synthetic receptors like chimeric antigen receptors (CAR) expressed on a T-cell cellular chassis has shown tremendous potential in cancer therapy. CD19 CAR T cells targeting the B-cell-specific antigen CD19 expressed in B-cell leukemia and lymphomas have shown durable responses that can last a decade^1–4^.

The clinical success of CAR T cells in turn has spurred extensive engineering of every element of the CAR, including the extracellular receptor, spacer, and intracellular domains^5,6^. Synthetic biology approaches have engineered synthetic receptors that allow the T cells to effectively discriminate antigen density or combinatorial antigens on target cells with the objective of maximizing efficacy while minimizing toxicity^7,8^. Similarly, the response of the T cells has also been successfully engineered to provide non-native functions, including the expression of additional receptors or the secretion of accessory anti-tumor proteins^9,10^. Despite these advances, the ability to take advantage of the synapse to facilitate the transport of non-cytotoxic proteins from T cells to the target cells has remained elusive. Engineered T cells with the ability to deliver on-demand non-cytotoxic cargo with specificity down to individual cells can advance our understanding of fundamental biological processes and expand our therapeutic options beyond the direct killing of tumor cells.

There are two major challenges in delivering non-cytotoxic passenger proteins through the immunological synapse. First, we do not understand the design rules for targeting proteins to the secretory lysosomes of T cells. The secretion of proteins with no directional specificity can be accomplished by appending an appropriate N-terminal leader peptide sequence; however, the mechanisms for sorting proteins into secretory lysosomes is complex and consequently a generalized targeting motif has not been identified^11,12^. The second challenge is that upon successful degranulation, the contents of the T cell lysosomes are delivered to the synapse and the passenger protein must be able to diffuse through perforin pores at the target cell membrane. There is an incomplete understanding of the features of passenger proteins (electrostatics, sterics etc.) that permit translocation through perforin pores.

Here, we demonstrate that the small, soluble, native lysosomal protein, NPC intracellular cholesterol transporter 2 (NPC2), can function as a non-cytotoxic lysosomal chaperone protein. By utilizing mCherry as the fluorescent reporter, we show that NPC2-mCherry fusion proteins are: packaged in the lysosome of T cells, are trafficked to the immunological synapse as part of secretory lysosomes, and can be detected in the target cells after delivery at the synapse. We demonstrate that NPC2 mediated delivery can be accomplished when activated through either the T-cell receptor (TCR) in mouse pmel CD8^+^ T cells or through a CAR (CD19-specific CAR) in both human CD4^+^ and CD8^+^ T cells. We propose that NPC2 is a lysosomal chaperone protein that can be used for the T cell mediated delivery of passenger proteins to target cells.

## Results

### Identification of NPC2 as a putative lysosomal chaperone

Our first objective was to identify an appropriate partner protein (chaperone) to enable the sorting of recombinant proteins into granules. The majority of lysosomal hydrolases, including granzymes, are modified with mannose 6-phosphate (M6P) residues^11^. Upon recognition of M6P residues, M6P-specific receptors (MPR) bind to the residues to form a clathrin-coated complex that can integrate with early endosomes, and the lysosomal protein gets released into the endosomes (**Fig. 1A**). We first investigated the use of granzyme B (GzB) as the lysosomal chaperone but unfortunately, the expression of granzyme B was toxic to T cells (not shown), and we focused on the identification of alternate non-cytotoxic proteins. In identifying alternate lysosomal chaperones, we focused on two specific design considerations: (1) the protein should be able to diffuse through perforin oligomers, and (2) the rate of diffusion through the perforin oligomers should be comparable to GzB.

**Figure 1.**
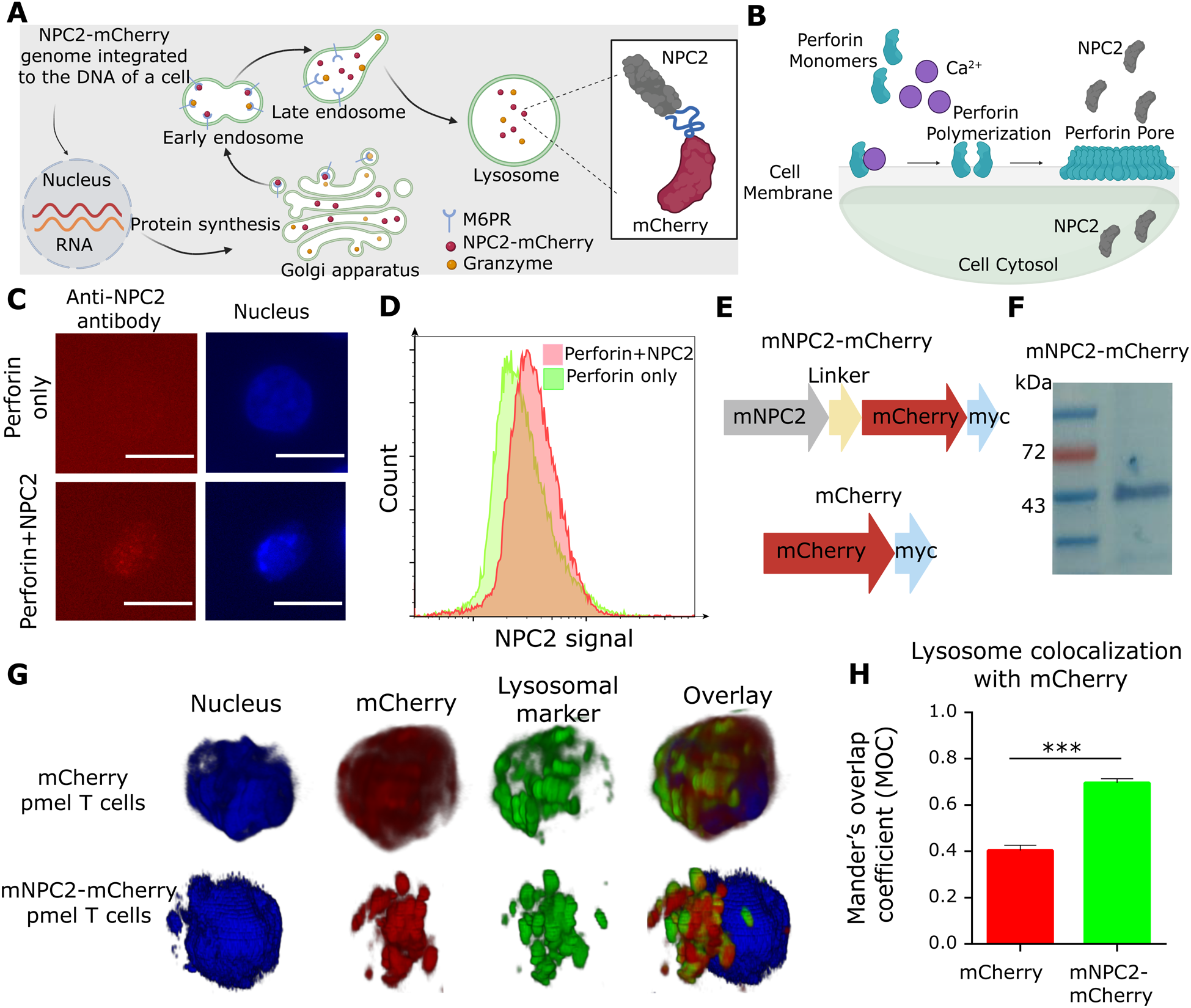
mNPC2 fusion proteins are localized to granules in mouse T cells. (A) Mechanism of sorting of proteins with M6P residues to granules. (B) Schematics of translocation of NPC2 protein through perforin pores. (C) Confocal images of the Jurkat cells incubated with purified perforin and NPC2 proteins. Scale bar, 10 μm. (D) NPC2 translocation was quantified for the Jurkat cells incubated with purified NPC2 and perforin proteins using flow cytometry. (E) Schematic of the constructs. mCherry was used to track the localization of mouse NPC intracellular cholesterol transporter 2 (mNPC2), and the linker was used to fuse the mNPC2 to the mCherry. (F) Western blot of the pmel T cells transduced with mNPC2-mCherry construct. (G) Subcellular localization of mNPC2 fused with mCherry in the granules of pmel T cells. (H) Manders’ coefficient quantified (MOC) colocalization of the mNPC2 and the lysosomal marker. The figure is plotted as Mean ± SEM (n = 200; t-test). *\*\*\*\*p < 0.0001; ***p < 0.001; **p < 0.01; *p < 0.05; ns: not significant.*

We chose NPC2 as a soluble candidate lysosomal chaperone protein based on the theoretical considerations associated with diffusion through perforin pores outlined below. The mechanism of protein translocation through perforin, and specifically the role of electrostatics, is actively debated^13,14^. Based on the crystal structure of human perforin (from PDB:5KWY), we mapped the electrostatic potential surface and projected this onto the known CryoEM map of oligomerized human perforin (**Figure S1A**). This map suggests that the top of perforin barrel (side closest to the T cell) harbors a positively charged surface and hence electrostatics is not a major determinant for the recruitment of passenger molecules to perforin pores (**Figure S1A**). Based on the map however, the lumen of the perforin barrel is negatively charged and hence can facilitate translocation of positively charged protein cargo (**Figure S1A**). Indeed, the structures of most granzymes presents a positively charged electrostatic surface (**Figure S1B**), and the pI of granzymes indicated a net positive charge at neutral pH (**Figure S1C**). Although the pI of NPC2 is 7.6 (positively charged in the lysosomes and likely neutral charge at the synapse), the protein’s surface has multiple positively charged amino acids that promote their electrostatic interaction with NPC1 or anionic phospholipids during cholesterol transfer (**Figure S2**)^15,16^. For these reasons, we hypothesized that from the perspective of electrostatics, NPC2 translocation through perforin would not be impeded. The second consideration for the transport across perforin is the delivery rate of the proteins being translocated. Delivery of lysosomal proteins to the target cytosol is a multi-step sequential process but the translocation across perforin pores is the rate-limiting step (detailed description of the model in the methods section). The rate of delivery of any lysosomal protein by perforin pores can be quantified by the rate constant k_g_ that is inversely proportional to radius of the molecule, r. Based on the size and structure of NPC2, we estimated the predicted radius of NPC2 to be ~2.23 nm which is smaller than the predicted radius of GzB (~2.75 nm). When NPC2 is used as a lysosomal chaperone, fusion proteins of sizes <32 kDa are expected to be translocated at a rate comparable to GzB. Thus, from a structural/biophysical perspective, we anticipated that NPC2 should function as a non-toxic lysosomal chaperone to facilitate delivery of proteins to the target cytosol.

The expression of proteins in T cells can be impacted by their differentiation status. Using single-cell RNA-seq data (dbGaP: phs002323.v1.p1), we compared the expression of *NPC2* across different subsets of CD8^+^ T cells. Unlike *GZB* expression that was heavily skewed towards the more differentiated CD8^+^ T cells, *NPC2* expression was uniform across all of the subsets of CD8^+^ T cells (naïve, central memory, effector memory and effector) [**Figure S1D**]. This suggested that the expression of NPC2 was not impacted by the differentiation state of the T-cell. Our next task was to directly test the translocation of the NPC2 protein across perforin pores. Accordingly, we incubated Jurkat cells with purified human perforin (hPerforin) and human NPC2 (hNPC2) proteins and detected the exogenously delivered hNPC2 using immunofluorescent confocal imaging and flow-cytometry (anti-NPC2 antibody) on fixed Jurkat cells. After 2 hours of incubation, cells treated with both hPerforin and hNPC2 showed higher intensity compared with controls (**Fig. 1C,D**). This result demonstrated that exogenous perforin is translocated through perforin pores and into the cytosol of target cells.

Since NPC2 is translocated through perforin pores, we next investigated its ability to act as a chaperone to facilitate the delivery of recombinant proteins to target cells. We evaluated the delivery capacity of NPC2 in three stages: (1) localization to granules in modified T cells, (2) trafficking of NPC2 containing secretory granules to the immunological synapse, and (3) delivery and translocation of NPC2 fusion proteins through perforin pores into the target cells.

### Recombinant NPC2 fusion proteins are sorted into mouse T cell granules

First, we investigated the expression of mouse NPC2-based fusion proteins and their localization in mouse T cells. To track the localization of NPC2 using live-cell microscopy, we constructed a genetic construct wherein the mouse NPC2 (mNPC2) gene is fused to mCherry from its C-terminal with a flexible hinge linker followed by a myc tag (**Fig. 1E**). We also separately cloned the mCherry gene into the same backbone, as a control (**Fig. 1E**). We generated retroviral particles and transduced transgenic T cells expressing the anti-gp100 TCR (CD8^+^ pmel T cells). Expression of mNPC2-mCherry had no adverse impact on the health of the pmel T cells, as evident from their normal proliferation rate. We confirmed that full-length mNPC2-mCherry was expressed by western blotting (**Fig. 1F**). We stained the T cells using Lysotracker deep red and quantified the localization of mCherry with respect to lysosomal granules (**Fig. 1G**). As expected by the restricted localization, pmel T cells expressing mNPC2-mCherry showed punctate mCherry staining, whereas pmel T cells expressing mCherry showed diffuse staining throughout the cytoplasm (**Fig. 1H**). Consistent with this observation, pmel T cells expressing mNPC2-mCherry showed significant localization of mCherry to lysosomal granules (Manders’ Overlap Coefficient, MOC, 0.70 ± 0.03) in contrast to pmel T cells expressing mCherry (MOC, 0.40 ± 0.06) [**Fig. 1I**]. This result illustrated that mNPC2 can function as a chaperone and promote the localization of fusion proteins to the lysosome of T cells.

### mNPC2-mCherry is localized in secretory granules that traffic to the immunological synapse in mouse T cells

Cytotoxic T lymphocytes (CTLs) mediate the killing of target cells by releasing lytic proteins that are stored in secretory granules at the immunological synapse (IS). Dynamic live-cell imaging of the granules can be used to monitor the localization of the secretory granules with respect to the IS (**Fig. 2A**). We incubated pmel T cells expressing mNPC2-mCherry or mCherry with MC38 target cells presenting gp100 (a tumor-associated antigen) and performed real-time confocal imaging. Within pmel T cells expressing mCherry, the mCherry signal was diffuse both before and after establishment of the synapse with no evidence of either punctate staining or specific trafficking to the IS (**Fig. 2B**). By contrast, pmel T cells expressing mNPC2-mCherry showed punctate staining prior to the formation of IS consistent with restricted localization to lysosomal granules (**Fig. 2B**). Upon establishment of the IS, the granules were localized to the contact area close to the IS and this localization is consistent with secretory granules that are trafficked to the IS before granule exocytosis permits release of the lysosomal cargo at the IS (Fig 2B). We quantified the distribution of the granules at the IS and this confirmed that the granules containing NPC2-mCherry is clustered at the IS (**Fig. 2C**). These results illustrate the mNPC2 fusion proteins are localized to secretory granules in T cells that converge at the IS upon recognition of tumor cells.

**Figure 2.**
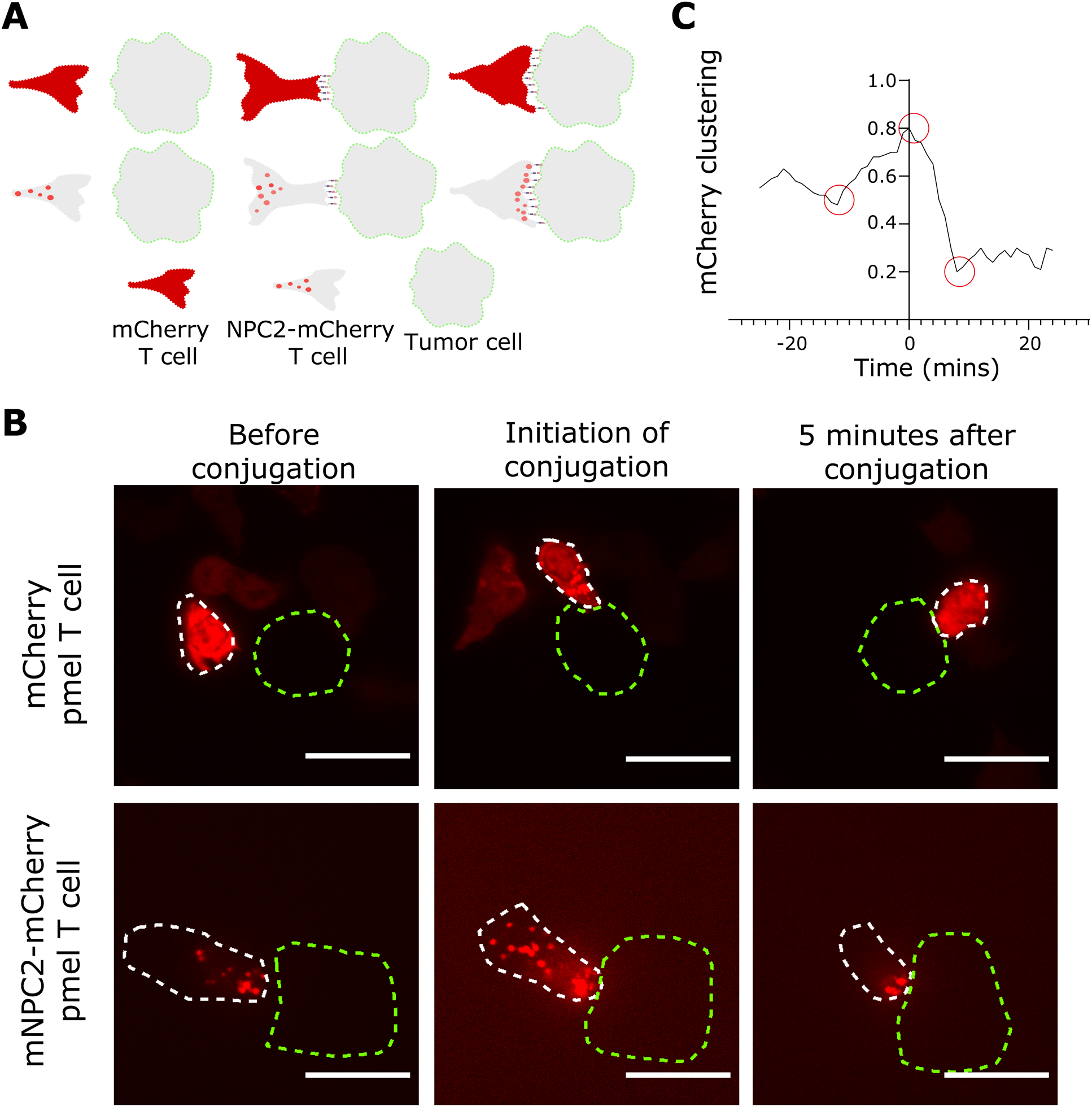
mNPC2-mCherry is sorted into secretory granules in T cells. (A) Schematic illustrating the localization of mCherry during stages of the T cell interacting with the tumor cell. (B) Representative image of the kinetics of lysosome trafficking recorded by confocal real-time imaging. For the sake of clarity, the outline of the pmel T cell and MC38/gp100 tumor cell are shown in white and green dotted dash lines, respectively. We analyzed a minimum of 60 events for both pmel-mCherry and pmel-NPC2-mCherry T cells. Scale bar, 10 μm. (C) Quantitative measurements of the distances between the T cell’s lysosomes before and after the conjugation with target cells as indicator of lysosomal polarization toward the synapse. The circles indicate the corresponding value for the pmel-mNPC2-mCherry in panel B.

### Mouse T cells can transfer mNPC2 fused proteins to the target cells

Since the live-cell imaging results suggested that mNPC2-mCherry is localized to the secretory granules in pmel T cells, we next investigated if we could directly quantify mNPC2-mCherry upon successful translocation and delivery into the target cells. We designed co-culture experiments to trace mCherry in target cells to test this hypothesis (**Fig. 3A**). Accordingly, we incubated labeled MC38/gp100 tumor cells with pmel T cells expressing either mNPC2-mCherry or mCherry for 2 hours and analyzed the cell populations using a flow cytometer. We gated live single cells corresponding to both the tumor cells and the pmel T cells (**Figure S4**) and measured mCherry fluorescence in these cells. Pmel T cells expressing mCherry exhibited higher fluorescence intensity compared to pmel T cells expressing mNPC2-mCherry (**Fig. 3B**). This was expected due to the smaller size and the cytoplasmic expression of mCherry in contrast to the lysosome localized expression of mNPC2-mCherry. Quantitative analysis of tumor cells demonstrated that the MC38/gp100 target cells co-cultured with pmel T cells expressing mNPC2-mCherry showed ~2.6-fold increase in the mCherry signal compared with target cells co-cultured with pmel T cells expressing mCherry (**Fig. 3C,D**). These results showed that despite being expressed at lower protein amounts per T cell, mNPC2-mCherry is more efficiently transferred to the tumor cells upon co-incubation in comparison to mCherry. This is consistent with an active role for mNPC2-mCherry trafficked and transferred at the IS.

**Figure 3.**
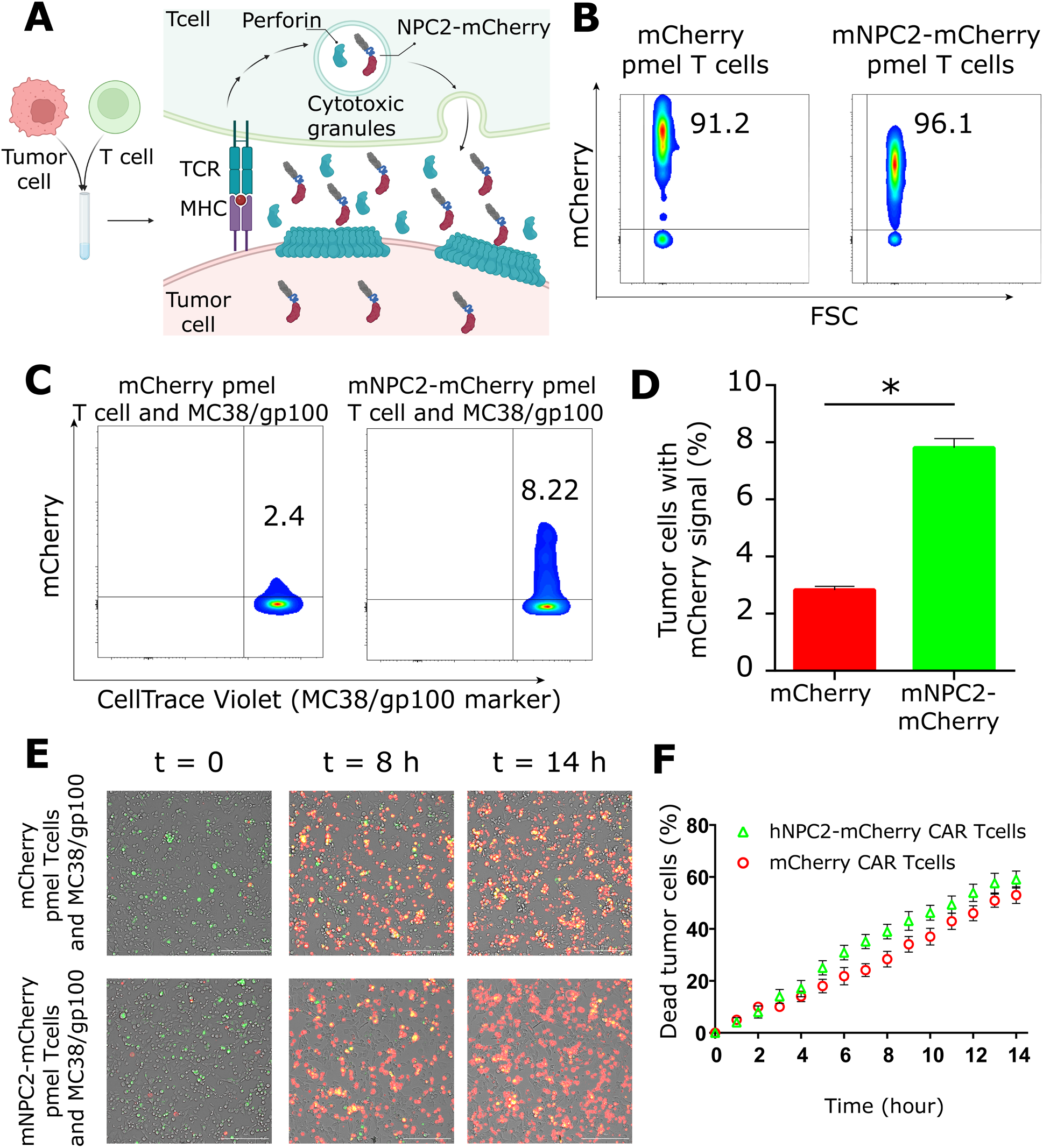
mNPC2-mCherry can be traced in the tumor cells and flow-cytometry. (A) Schematics of co-culture assay. Upon conjugation of T cells and tumor cells, the content of granules within NPC2-mCherry transduced tumor cells are released into the synapse. Perforin monomers bind to the target cell membrane and oligomerize to form perforin pores. Formation of the pore enables delivery of NPC2-mCherry fusion protein into the cytosol of tumor cells. (B) Flow cytometry plots show the distribution of the mCherry in transduced T cells. (C) Tumor cells were assayed for transferring of mNPC2-mCherry through the immunological synapse to the tumor cells after 2 hours of incubation. (D) Percentage of the mCherry in MC38/gp100 tumor cells after co-culture assay plotted as Mean ± SEM (n = 4; t-test). (E) Representative images of tumor cells incubated with pmel T cells at different time points. Green indicates labelled tumor cells and red indicates dead cells. Scale bar, 50 μm. (F) Pmel T cells transduced with mNPC2-mCherry killed the MC38/gp100 tumor cells at the same rate as mCherry pmel T cells. *\*\*\*\*p < 0.0001; ***p < 0.001; **p < 0.01; *p < 0.05; ns: not significant.*

Overexpression of essential proteins like mNPC2 that are targeted to restricted organelles like the granules can impede cellular function. To evaluate whether the mNPC2 overexpression can impact cytotoxicity of the T cells, we measured the number of MC38/gp100 targets cells killed by pmel T cells using dynamic live cell imaging (**Fig. 3E**). Tracking the tumor cells during 15 hours of co-incubation with pmel T cells revealed a similar killing rate for T cells expressing mNPC2-mCherry and T cells expressing mCherry cells (**Fig. 3F**). Collectively, these data suggest that mNPC2 fused proteins can be translocated into the tumor cells at the IS without adversely impacting T cell killing.

### hNPC2 targets mCherry to secretory granules in human T cells

To investigate if hNPC2 can function as a lysosomal chaperone in human T cells, we cloned the human NPC2 (hNPC2) downstream of the CD19 chimeric antigen receptor (CAR) gene separated by a self-cleaving peptide, T2A, to enable the expression of both proteins (**Fig. 4A**). We separately cloned a construct containing the CAR gene followed by mCherry (no NPC2) as a control (**Fig. 4A**). We generated retroviral particles and manufactured CAR T cells by our standard 10-day expansion protocol^17^. Phenotyping the cells showed that the majority of T cells expressing both CAR constructs were memory T cells (CD45RO^+^, **Fig. 4B**). Like the mouse T cell data, human CAR T cells expressing NPC2-mCherry showed punctate distribution with significant localization to the lysosomal granules (MOC, 0.63 ± 0.05) compared to human CAR T cells expressing mCherry (**Fig. 4C,D**). We also compared the cytotoxicity of the CAR T cells against the NALM6 tumor cells with dynamic live-cell imaging. NALM6 killing results showed a similar rate for both NPC2-mCherry and control CAR T cell groups (**Fig. 4E**).

**Figure 4.**
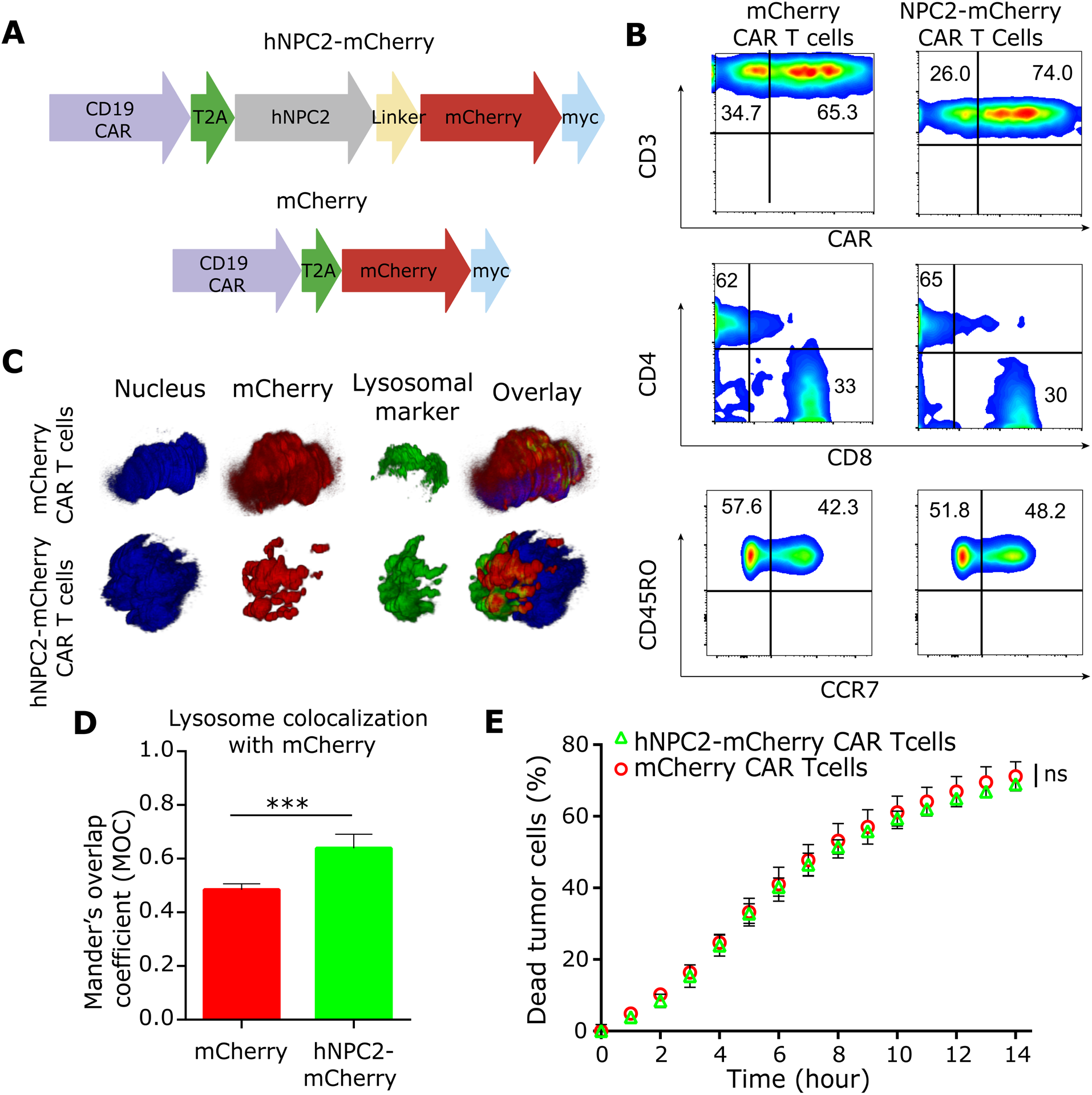
Expression and tracing of proteins targeted to granules within human T cells. (A) Schematic of the constructs. mCherry was used to track the localization of human NPC intracellular cholesterol transporter 2 (NPC2), and the linker was used to fuse the mouse NPC2 (mNPC2) to the mCherry. CD19 CAR was added to the N terminus. (B) Phenotyping of the CAR T cells. (C) Subcellular localization of hNPC2 fused with mCherry and mCherry alone in the granules of the CAR T cells. (D) Colocalization of the hNPC2 and the lysosomal marker was quantified by Manders’ coefficient. The graph is plotted as Mean ± SEM (n= 4; t-test). (E) CAR T cells transduced with hNPC2-mCherry killed the NALM6 tumor cells at the same rate as mCherry CAR T cells. *\*\*\*\*p < 0.0001; ***p < 0.001; **p < 0.01; *p < 0.05; ns: not significant.*

### Overexpression of hNPC2-mCherry does not impair the function of human T cells

We next used live-cell imaging to evaluate the trafficking of mCherry upon the interaction of the human CAR T cells with CD19 expressing tumor cells, NALM6. To quantify differences in phenotypes (CD4^+^ and CD8^+^ T cells), we evaluated each of these subsets separately within hNPC2-mCherry and mCherry expressing CAR T cells. Consistent with localization to secretory granules, both CD4^+^ and CD8^+^ T cells expressing hNPC2-mCherry showed punctate staining before the formation of the IS (**Fig. 5A)**. Similar to the mouse CD8^+^ T cells, upon formation of the IS, both human CD4^+^ and CD8^+^ T cells showed clustered and localized expression of NPC2-mCherry at the contact area close to the synapse (**Fig. 5A**). These results confirmed that NPC2 functions as a lysosomal chaperone to target mCherry to secretory granules in both CD4^+^ and CD8^+^ human T cells.

**Figure 5.**
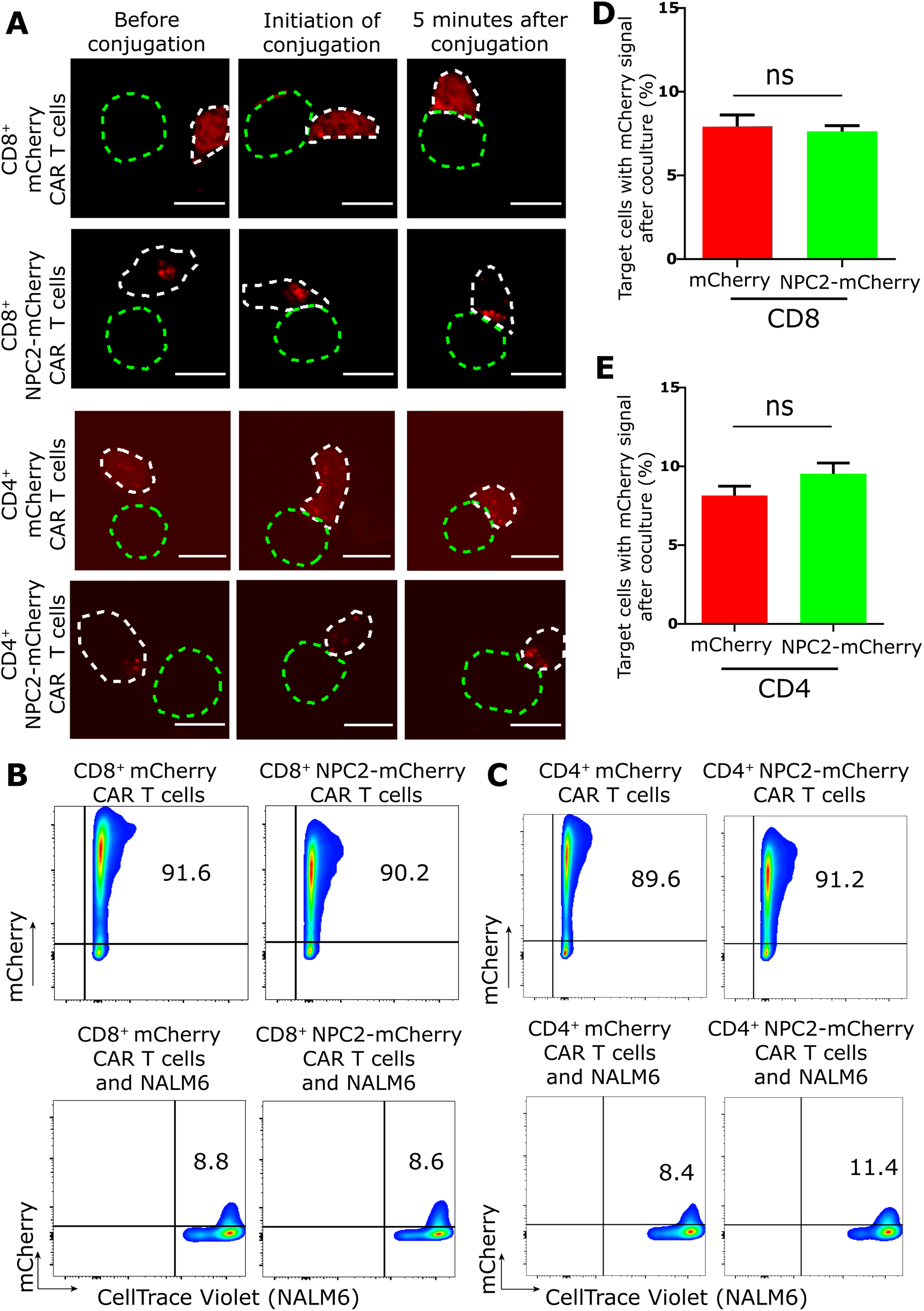
CAR T cells can initiate T cell killing steps and transfer the hNPC2-mCherry to the NALM6. (A) Representative image of the kinetics of lysosome trafficking recorded by confocal real-time imaging. For the sake of clarity, the outline of the CAR T cell and NALM6 tumor cell are shown in white and green dotted dash lines, respectively. We analyzed a minimum of 60 events for both mCherry and NPC2-mCherry CAR T cells. Scale bar, 10 μm. (B,C) Tumor cells were assayed for transferring of hNPC2-mCherry by CD8^+^ or CD4^+^ T cells through the immunological synapse to the tumor cells after 2 hours of incubation. (D,E) Percentage of the mCherry in NALM6 tumor cells after co-culture with CD8^+^ or CD4^+^ T cells plotted as Mean ± SEM (n = 4; t-test). *\*\*\*\*p < 0.0001; ***p < 0.001; **p < 0.01; *p < 0.05; ns: not significant*.

We next sought to test the efficacy of translocation for human NPC2 fusion from CAR T cells into the human tumor cells. We designed a co-culture experiment similar to the one performed with the pmel T cells. We labeled the NALM6 tumor cells and co-incubated them with either CD4^+^ or CD8^+^ CAR T cells (**Fig. 5B,C**). While both CD4^+^ and CD8^+^ T cells were able to transfer NPC2-mCherry to tumor cells, this was no different from these subsets of cells expressing mCherry alone (**Fig. 5D,E**). Careful comparison of the mouse and human T cells suggests that the primary difference arises in mCherry transfer without the NPC2 chaperone (**Fig. 3D and 5D,E**). We posit that the elevated expression of mCherry relative to NPC2-mCherry (**Fig. 5B,C**), the efficiency of synapse formation with the CAR in human T cells, and the known propensity for trogocytosis to permit mCherry transfer in CAR T cells all contribute to the elevated transfer of mCherry in the absence of NPC2^18^.

## Discussion

T cells have the ability to recognize and respond to even a single copy of their cognate peptide displayed on the target cell, making them very powerful sentinels that patrol the human body. This ability to identify and eliminate single cells with exquisite specificity has enabled their development as living drugs. The engineering and expression of CARs within T cells has expanded the targeting capability of T cells and has accelerated their clinical application. Despite these advances, the cellular response upon activation of the TCR/CAR has primarily focused on cytotoxic responses leading to the death of the target cells. The ability to deliver non-cytotoxic cargo via the IS can expand the range of biotechnological and even clinical application of T cells.

The trafficking of recombinant proteins via the lysosomal pathway is unfortunately not straightforward. The sorting of lysosomal proteins into granules can be either M6P receptor dependent or M6P receptor independent. M6P phosphorylated proteins interact with two different types of receptors: cation-independent and cation-dependent mannose 6-phosphate receptors^19,20^. Mass spectrometry analysis have revealed multiple randomly distributed M6P sites on soluble lysosomal proteins^21^. These sites are distinguished by their specific conformations that are recognized by multiple enzymes selectively phosphorylating N-linked high mannose oligosaccharides in Golgi compartments^22^. These recognition motifs can be distributed throughout the protein and consequently there is no singular peptide motif that has been identified that can permit the trafficking of recombinant proteins to lysosomes. Although M6P receptors likely play a dominant role in the sorting of lysosomal proteins, at least two alternative receptors have been identified: the lysosomal integral membrane protein (LIMP-2) and sortilin^23–25^. Unfortunately however, the rules for how sorting is accomplished through these alternate receptors is even less well-characterized^25^. In the absence of defined targeting/trafficking motifs or peptides, the most common well-utilized mechanism to traffic recombinant proteins to the lysosomes is via fusion to native lysosomal proteins. We chose NPC2, a native intralysosomal protein as a lysosomal chaperone in T cells for several reasons outlined below.

NPC2 displays biophysical characteristics making it an attractive candidate as a lysosomal chaperone. First, NPC2 is a globular protein with a comparatively small size (16 kDa), which can be advantageous for its application in the expression and delivery of fusion proteins. Viral transduction has been shown to be an efficient and safe tool for the expression of proteins in T cells. Previous studies have revealed the impact of genome size on producer cell mRNA levels, packaging efficiency, and infectivity of the virions^26^. Therefore, NPC2’s small size can be beneficial when engineering virions for the delivery of synthetic receptors. Second, the small size of NPC2 coupled with its elongated shape and surface positive charges make it an ideal partner to facilitate translocation through perforin pores (16nm in diameter)^27^. The overexpression of NPC2 is also likely to have beneficial effects to T cells. The native function of NPC2 is to facilitate the export of cholesterol out of the lysosomes by enabling the transfer of the cholesterol to the sterol-binding pocket of NPC intracellular cholesterol transporter 1 (NPC1)^28^. The NPC2-NPC1 axis thus modulates the total amount of free cholesterol in cells. An increase in the plasma membrane cholesterol level of T cells increases membrane fluidity making the T cells more responsive to TCR signaling and an enhanced ability to form stable IS^29^.

Our work demonstrates the critical first step in the delivery of non-cytotoxic proteins via T cells using NPC2 as the lysosomal chaperone. In designing our experiments, we used mCherry as the payload for delivery. Several design considerations will limit the payloads that can be delivered using the NPC2 system including: (1) size, charge, and structure of the protein payload; and its ability to be transported through perforin pores, (2) stability of the payload in the acidic environment of the lysosomes, and (3) the ability of payloads to tolerate fusion at its N-terminus. Fortunately, there are multiple classes of thermostable proteins that satisfy these criteria and hence provide exciting opportunities for the delivery of non-cytotoxic payloads to modify target cells.

Broadly, we have combined the granzyme-perforin pathway with a non-cytotoxic protein for cell-to-cell delivery of fusion proteins. Inserting fusion therapeutic proteins into T cell-mediated killing can be a powerful approach to improving the efficacy of engineered T cells by synthetic biology.

## Acknowledgments

This publication was supported by the NIH (R01GM143243), CPRIT (RP180466); MRA Established Investigator Award to NV (509800), Welch Foundation (E1774); NSF (1705464); CDMRP (CA160591); Owens foundation; Cytometry and Cell Sorting Core at Baylor College of Medicine with funding from the CPRIT Core Facility Support Award (CPRIT-RP180672), the NIH (P30 CA125123 and S10 RR024574) and the expert assistance of Joel M. Sederstrom.

## Author contributions

N.V. conceived this study. N.V. and A.S. designed the study. A.S., I.B., A.L., M.M., and R.B. performed the experiments. N.V. and A.S. wrote the paper. A.S., M.F., K.L.W, A.R, W.P., and N.V. analyzed the data. All authors discussed the results and commented on the manuscript.

## Materials and Methods

### Cell culture

Platinum (plat-E and plat-GP) retroviral packaging cell lines and splenocytes were cultured in DMEM (Gibco cat. # 11995065) and MEM α (Gibco cat. # 12571089) media, respectively. MC38/gp100 murine colon adenocarcinoma, NALM6 cells, and PBMCs were cultured in RPMI 1640 medium. All the media were supplemented with 10 % (v/v) fetal bovine serum (R&D systems cat. # S11550), 1 % (v/v) HEPES (Fisher cat. # 15630106), 1 % (v/v) Glutamax 100X (Fisher cat. # 35-050-061), and 50mg/ml Normocin (InvivoGen cat. # ant-nr-2).

### Construction of vectors

Murine *NPC2* was amplified from Integrated DNA Technologies (IDT) GeneBlock. It has been previously shown that 11 N terminal residues of mCherry are prone to cleavage by the lysosomal proteases^30^. Therefore, we deleted these residues from the construct design. A flexible linker (EFPKPSTPPGSSGGAP) that can span 2.5-2.7 nm^31^ was used to create a gap between mNPC2 and mCherry in the fusion protein. *Myc-tag* sequence was added to the C terminal to detect the proteins’ expression by western blotting. All the genes were subcloned into an RVKM vector (gift from Dr. Weiyi Peng). Sequences of all constructs were verified. For cloning human *NPC2* gene, we prepared cDNA of the U2OS cell line because of its high expression level of NPC2. Next, we cloned the *NPC2, mCherry,* and *myc* genes into a retroviral vector (containing *CD19 CAR* gene) using NEB HiFi DNA Assembly Master Mix (NEB cat. # E2621S).

### *In Vitro* translocation assay

Purified human perforin was purchased from Raybiotech (cat. # 230-00687). Purified human NPC2 was purchased from Acrobiosystem (cat. # NP2-H52H1). The sublytic concentration of perforin was determined with a flow cytometer (15 μg/ml). The jurkat cells were equilibrated in a solution of 0.4% Bovine serum albumin (BSA), 2 M CaCl2, and 1% HEPES for 15 minutes before addition of perforin and NPC2 proteins. The cells were incubated with the proteins for two hours and washed twice after the incubation. The images of the cells were captured as explained below. The samples were analyzed with a BD LSRFortessa. The flow cytometry data was analyzed with FlowJo software version 10.8 (Tree Star Inc, Ashland, OR, USA).

### Mathematical modeling of translocation through perforin pores

It has been previously shown that a finding the perforin pores by lysosomal proteins is a rate-limiting step and occurs at the rate of k_g_=3πR_p_^2^D/l^2^h^2^, in which R_p_, l, and h are constant for all proteins and D is the diffusivity of the protein^32^. We can estimate the diffusivity of proteins by the Stokes-Einstein relation, D_free_ = k_B_T/(6πη_w_r), where k_B_T is the thermal energy, ⊓w is the solvent viscosity, and r is the radius of the molecule^32^. Knowing that all the molecules are in the same condition in a solution, we can argue that k_g_ is inversely correlated to the radius of a molecule, r.

### Retroviral transduction of pmel T cells

Pmel T cells were transduced using the protocol previously described^33^. In brief, we produced retroviral particles by transfecting plat-E cells with the retroviral vectors and packaging plasmids using lipofectamine 2000 transfection reagent (Invitrogen cat. # 11668-027). Viral particles were collected after 48 h. Splenocytes were harvested from the spleen of pmel-1 mice and activated with 500 U/ml hIL-2 (Proleukin, Bayer Healthcare Pharmaceuticals), 50 mM 2-mercaptoethanol (Fisher cat. # 31350010), and 0.5 μg/ml anti-mouse CD3 (BD Bioscience cat. # 553057). After 24 h, the viral particles were concentrated using Amicon^®^ Ultra-15 Centrifugal Filter Unit (Millipore cat. # UFC910024). The activated splenocytes were infected by adding the concentrated virus and 1.6 μg/ml Polybrene and spinning the plate at 2000 rpm for 90 minutes. The T cells were propagated for three days and sorted with BD FACS Melody Cell Sorter.

### Retroviral transduction of PBMCs and phenotyping

RD114 expressing PlatGP cells were transfected with the constructs using Lipofectamine LTX Reagent with PLUS Reagent (Thermofisher cat. # A12621). For activation of PBMCs, we coated a non-treated 24-well plate with an anti-CD3 antibody overnight. The next day, we seeded 1.5×10^6^ PBMCs resuspended in 1 mL of RPMI 1640 media (10% FBS), supplemented with anti-CD28 antibody, IL-7 (Peprotech cat# 200-07, 10 ng/mL), and IL-15 (Peprotech cat# 200-15, 5 ng/mL). After 48 hours, the activated T cells were transduced with retroviral supernatants by centrifugation onto a retronectin-coated plate. New media supplemented with IL-7 and IL-15 was subsequently added to the cells every 2 to 3 days. For phenotyping, CAR T cells were stained for 30 min at 4°C using a panel of human-specific antibodies from BD Biosciences: CD3 (SK7), CD4 (OKT4), CD8 (RPA-T8). In addition, cells were co-stained with the viability dye 7-AAD (BD) and the in-house anti-CD19scFv.

### Confocal microscopy

For live-cell imaging, transduced T cells were labeled for granules by incubating with LysoTracker Deep Red in media to a final concentration of 100 nM at 37°C for 1 hour. The nucleus was stained with Hoechst 33342 (Sigma, 14533, 10 μg/ml) for 20 min at 37°C and washed twice with PBS before acquiring the images. The unlabeled tumor cells and transduced T cells were added to a 96 Well Black Plate glass bottom plate (Thermo Scientific, 160376). 60-70 z-stacks (0.2μm steps) were captured by a Nikon (Minato, Tokyo, Japan) Eclipse Ti2 inverted microscope equipped with a 100x, Nikon, Plan Apo Lambda, oil, 1.45 NA objective from different fields of view using DAPI, TXRed, and Cy5 channels at 1 min intervals for 1 hour. The composite images were created using NIS-Elements Viewer software.

### Analysis of confocal images

We extracted z-stacks of 16-bit images for each channel and processed them in ImageJ (National Institutes of Health (NIH), USA) using a series of plugins. First, we applied 3D watershed, 3D objective counter plugin, and 3D ROI Manager plugin to the channels corresponding to mCherry and LysoTracker Deep Red. Then, we obtained the pixel values for each voxel of detected mCherry or Lysosomal marker objects. Next, we used nucleus staining to detect single cells by obtaining their location in the image. Finally, we analyzed all the processed outputs in R program to calculate the Mender’s colocalization coefficient between mCherry and lysosomal marker objects for every single cell.

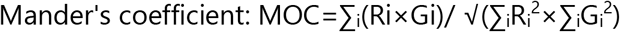

We quantified the clustering of lysosomes through blob detection using Python 3.7 with scikit-image library 0.19.2. For each image taken by the confocal microscope, we applied blob detection with Laplacian of Gaussian as kernel to detect the mCherry signal blobs. We only accept detections with a radius between 2-7 pixels and a sigma value of Gaussian higher than 0.03. After detection, we calculated the Euclidean distance between both centroids of each pair of blobs, and their variance is the lysosome clustering value.

### Co-culture Experiment

We labeled MC38/gp100 and NALM-6 cells with Cell-Trace Violet (Invitrogen cat. # C34557) dye at a concentration of 5 μM. The MC38/gp100 cells were loaded into a 6-well plate until they formed their spindle-like morphology. The NALM6 cells were seeded in a round-bottom plate. The T cells were incubated with the cancer cells at a 5:1 effector:target (E:T) ratio. After 2 hours, the media was collected into a round-bottom polystyrene test tube. The cells were washed twice with PBS and resuspended in FACS buffer containing Sytox green (Invitrogen cat. # S34860) at a final concentration of 100 nM to distinguish the live and dead cells. The samples were analyzed with a BD LSRFortessa. The flow cytometry data was analyzed with FlowJo software version 10.8 (Tree Star Inc, Ashland, OR, USA).

### Cytotoxicity assay

For real-time cytotoxicity assay, we used Cytation 7 Cell Imaging system that allows us to monitor T cell killing over time quickly. CellVue (MilliporeSigma cat. # MINCLARET-1KT) labeled MC38/gp100 cancer cells, and the transduced pmel Tcells were resuspended in MEM media containing Sytox green as death marker. They were incubated in a 96 well-plate at 1:1 effector:target (E:T) ratio under 37°C and 5% CO2. The images were captured by Cytation 7 inverted microscope using 20x Plan Fluorite WD 6.6 NA 0.45 objective from FITC and TXRed channels at 1 hour timeframe for 24 hours. The primary mask was automatically applied to the images based on the Cy5 channel in Gen5 software. The detected target cells with an intensity unit of higher than 500 were counted as cells killed by pmel T cells. The number of dead cells over time was plotted in Prism software.

### Western blotting

Transduced pmel T cells (1 x 10^6^) were lysed in radioimmunoprecipitation assay (RIPA) buffer (2 mM ethylenediaminetetraacetic acid (EDTA), 150 mM NaCl, 50 mM Tris-HCl, and 1% Triton X-100) containing protease inhibitors and phosphatase inhibitors and spun down for 20 min at 13000rpm at 4 °C. The supernatants were quantified for protein concentration using BCA Protein Assay (Pierce, Thermo Fisher Scientific). The protein samples were separated on 4–15% Mini-PROTEAN^®^ TGX™ Precast Protein Gels (Biorad cat. # 4561086) and transferred to a Hybond Amersham PVDF transfer membrane (MilliporeSigma cat. # GE10600023 and blocked with 5% skimmed milk in TBST for 1 hour at RT. The PVDF membrane was incubated with the anti-c-myc primary antibody (Biolegend; Clone 9E10) diluted in 2.5% bovine serum albumin (BSA) (1:1000) and kept at 4 °C overnight. The membrane was washed with TBST and then incubated with horseradish peroxidase (HRP)-conjugated secondary antibody at dilution of 1:3000 diluted with 2.5% BSA (Cell Signaling Technology cat. # 7076). The protein expression was visualized by using TMB blotting solution.

**Supplementary Figure 1.**
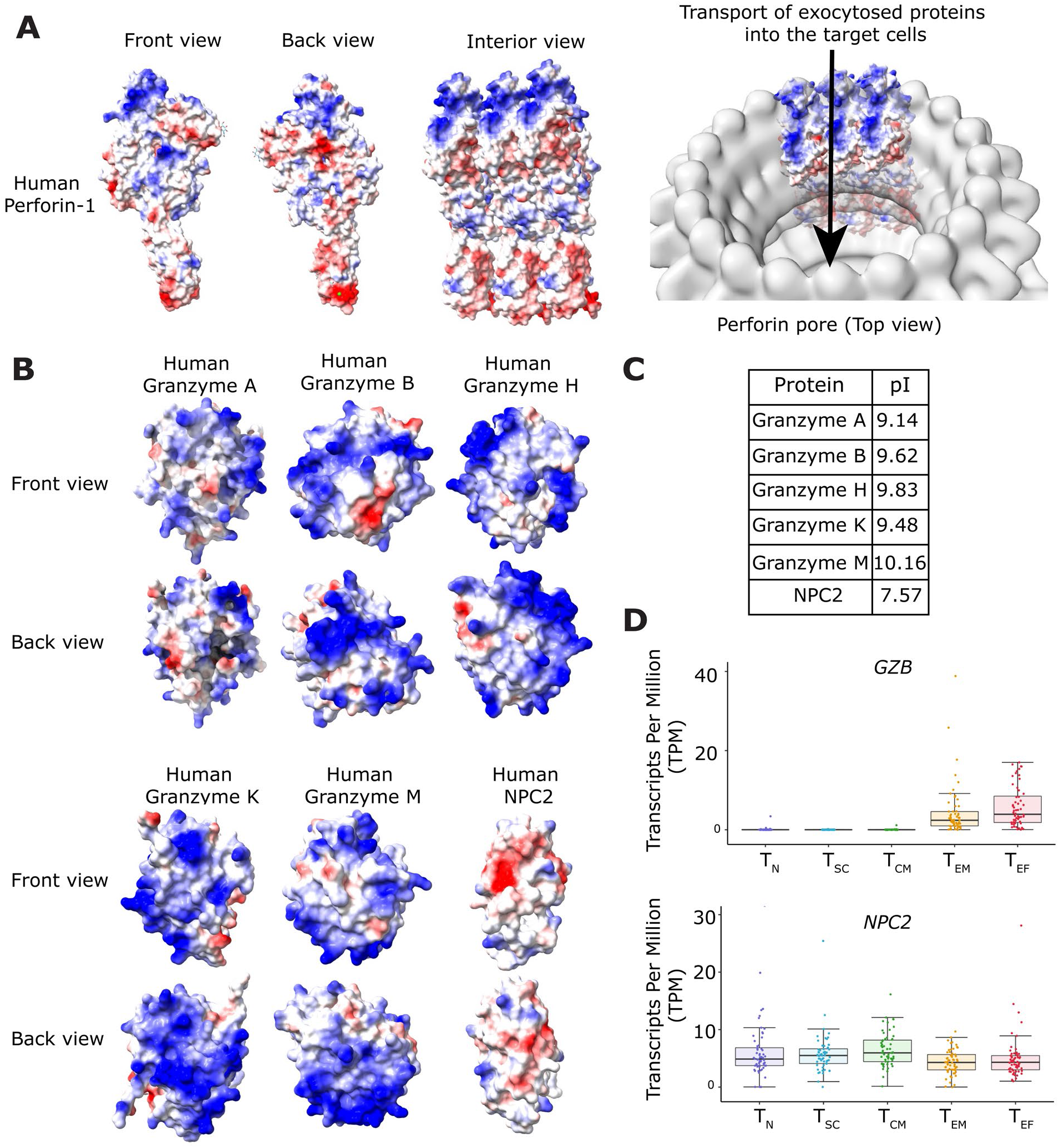
(A) Electrostatic surface potential map of Perforin-1 monomer (PDB: 3NSJ) and electron microscopy map of perforin pore (Accession number: EMD-1769). (B) Electrostatic surface potential maps of Granzyme A (PDB: 1OP8), Granzyme B (PDB: 1FQ3), Granzyme M (PDB: 2ZGC), Granzyme H (PDB: 3TJU), and NPC2 (from PDB: 5KWY). Blue indicates positive charges and red indicates negative charges. (C) PI of the proteins in Fig. S1B. (D) Expression of *GZB* and *NPC2* within different subsets of T cells (based on dbGaP: phs002323.v1.p1).

**Supplementary Figure 2.**
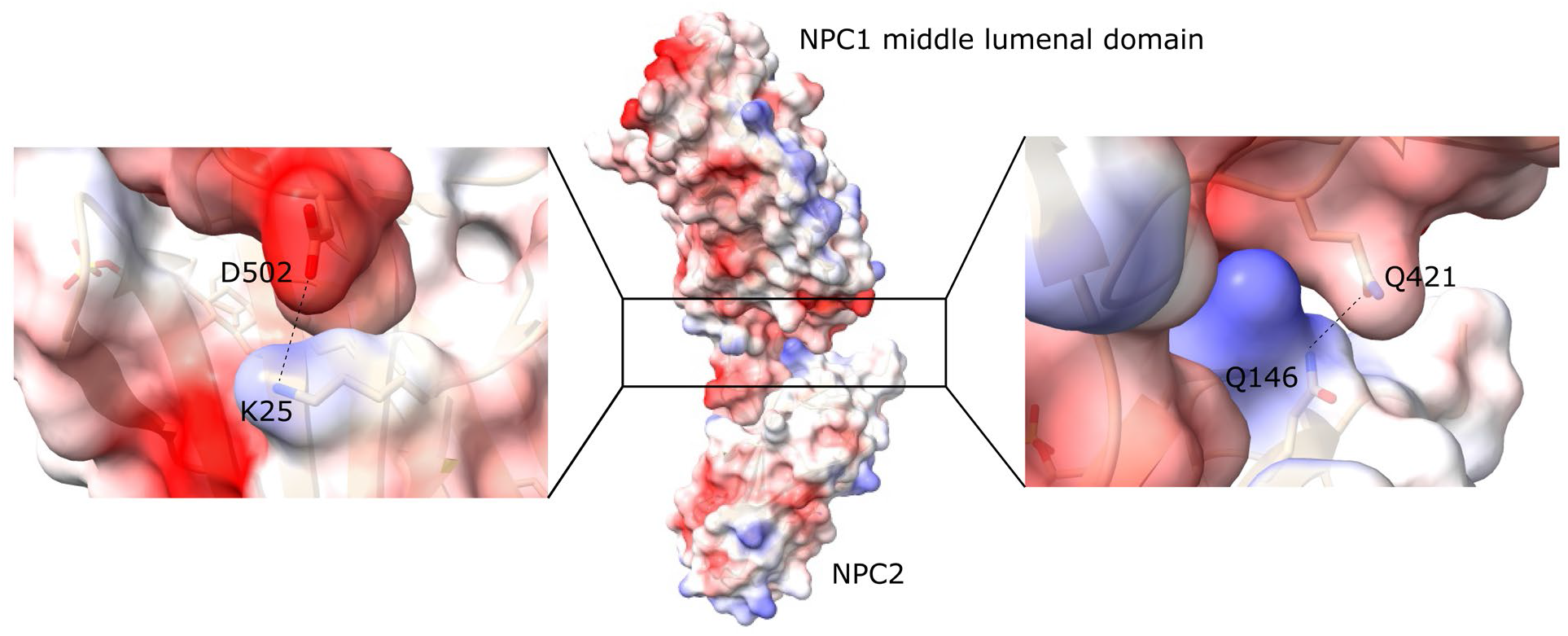
Interaction between positively charged amino acids (shown as blue) on NPC2 protein and negatively charged amino acids on NPC1 protein (shown as red) (PDB: 5KWY)

**Supplementary Figure 3.**
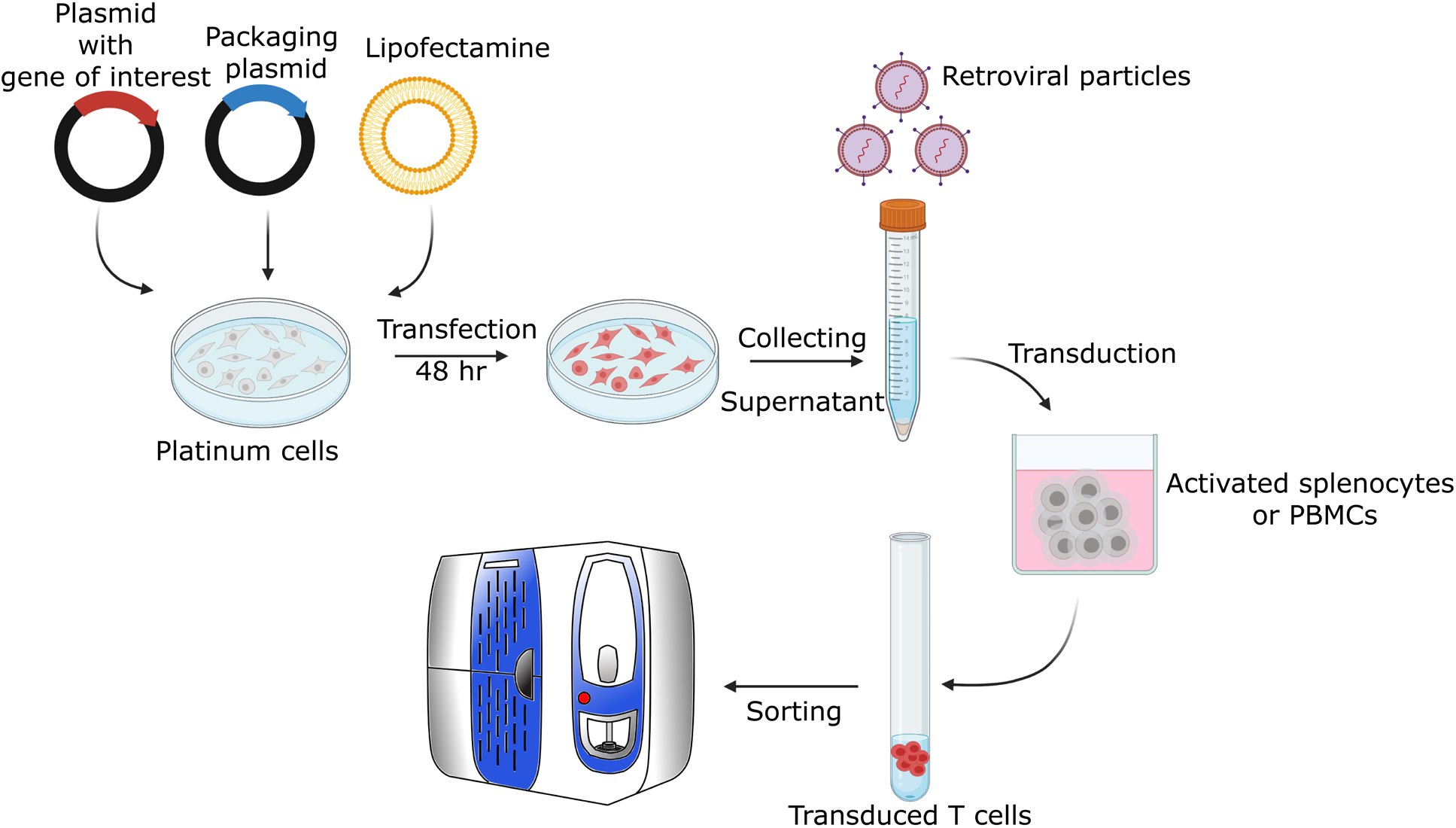
Schematics of the retroviral transduction of mouse T cells.

**Supplementary Figure 4.**
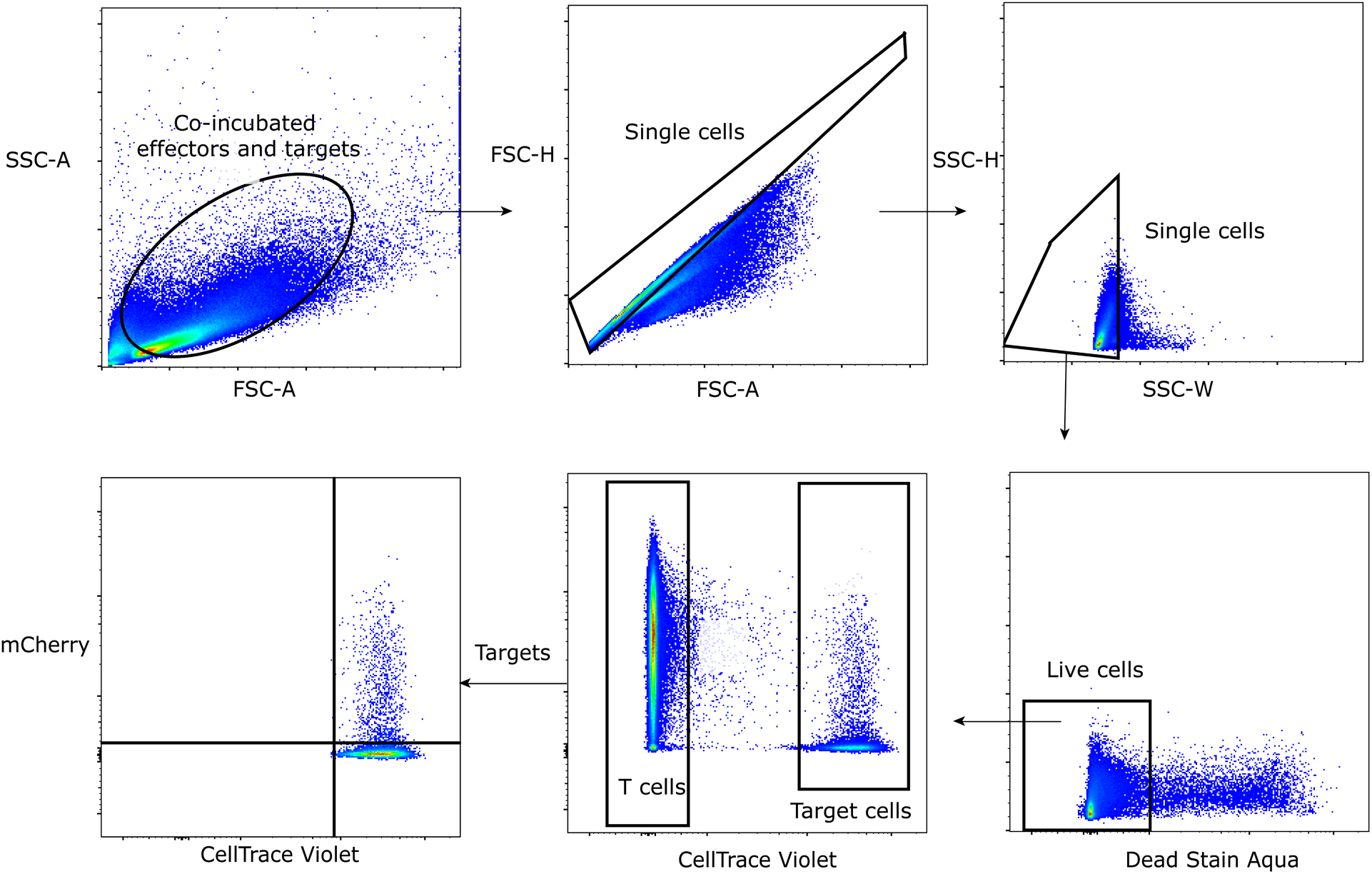
Gating strategy for analyzing the translocation of NPC2-mCherry to the tumor cells.

**Supplementary Figure 5.**
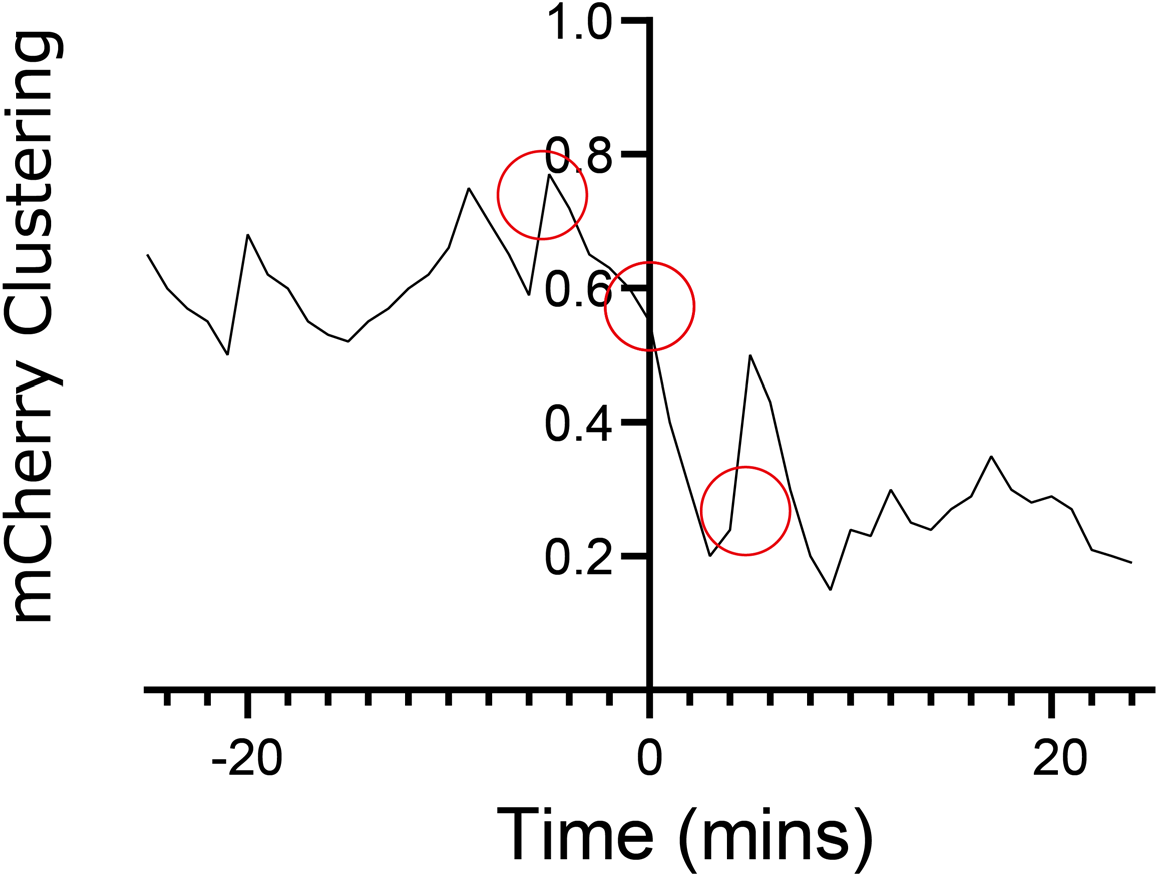
Quantitative measurements of the distances between the T cell’s lysosomes before and after the conjugation with target cells as indicator of lysosomal polarization toward the synapse. The circles indicate the corresponding value for the hNPC2-mCherry CAR T cells in Fig. 5A, row2.

